# Reproducibility of importance extraction methods in neural network based fMRI classification

**DOI:** 10.1101/197277

**Authors:** Athanasios Gotsopoulos, Heini Saarimäki, Enrico Glerean, Iiro P. Jääskeläinen, Mikko Sams, Lauri Nummenmaa, Jouko Lampinen

## Abstract

Recent advances in machine learning allow faster training, improved performance and increased interpretability of classification techniques. Consequently, their application in neuroscience is rapidly increasing. While classification approaches have proved useful in functional magnetic resonance imaging (fMRI) studies, there are concerns regarding extraction, reproducibility and visualization of brain regions that contribute most significantly to the classification. We addressed these issues using an fMRI classification scheme based on neural networks and compared a set of methods for extraction of category-related voxel importances in three simulated and two empirical datasets. The simulation data revealed that the proposed scheme successfully detects spatially distributed and overlapping activation patterns upon successful classification. Application of the proposed classification scheme to two previously published empirical fMRI datasets revealed robust importance maps that extensively overlap with univariate maps but also provide complementary information. We conclude that importance maps are superior to univariate approaches for both detection of overlapping patterns and patterns with weak univariate information.

## 1. Introduction

Multivariate pattern analysis (MVPA) has been established as an indispensable tool for fMRI research since its introduction by Haxby and colleagues in 2001. It has been shown to overcome limitations of univariate methods by addressing voxel activation collectively in terms of distributed patterns (Norman et al., 2006; Lewis & Peacock, 2013; Cohen et al., 2017) and thus has emerged as a powerful analytic technique in both experimental and clinical settings. MVPA typically refers to a set of machine learning methods, applicable to fMRI data, that collectively analyze voxel activity.

Classification, as a subset of MVPA methods, aims at establishing discriminability between conditions such as brain activity patterns elicited by seeing different object categories. Studies in fMRI classification are often confined in reporting classification accuracy, which is an informative measure with direct impact in clinical diagnosis tools (Coutanche et al., 2011; Sundermann et al., 2014). However, to gain a better understanding of information representation in the brain, it is equally important to identify which regions drive the classification especially in whole brain inter-subject classification.

This is a general problem in core machine learning and image classification (Montavon et al., 2017). Main goal of these techniques is to extract meaningful and consistent patterns that represent the decision boundary of the classifier, or in other words, patterns that lead the classifier to a particular decision. In the case of image classification, these patterns refer to pixels of the image while in fMRI classification they typically refer to voxels. Evaluation of such methods is more intuitive in image classification where visual inspection is a safe option, but interpretation of brain activity patterns in a three dimensional space is far from trivial. Such patterns in fMRI classification have been addressed by previous studies and are often referred to as importance maps (Polyn et al., 2005), relevance maps (Åberg & Wessberg, 2008; Schrouff & Phillips, 2012) or sensitivity maps (Rasmussen et al., 2011) but there is yet no rule of thumb for their extraction.

In linear classifiers, a typical approach is to visualize the weights (Pereira et al., 2009), or the weights-input product (Polyn et al., 2005); this is however not feasible for nonlinear models such as kernel based models (Rasmussen et al., 2011) or deep neural networks. There has been no general proof for superiority of non-linear classifiers over linear classifiers in fMRI data analysis (Haxby et al., 2014; Kamitani & Tong, 2005; Misaki et al., 2010), although there are hints that such nonlinearities do exist as co-activation of two or more brain regions may be necessary to trigger certain neural mechanisms (Kober et al., 2008).

Failure to detect such nonlinearities is partly attributed to the “curse of dimensionality” (Cohen et al., 2017) where the number of parameters to be trained is much higher than the number of samples. This often leads to overfitting, that is, low generalization of the model. The problem of dimensionality is commonly tackled by reducing the number of voxels involved in the classification either by performing a univariate feature selection step prior to classification (Coutanche et al., 2011; Kohler et al., 2013; Sitaram et al., 2011) or by performing localized analyses. Main drawback of the former approach is that it might remove voxels that contain multivariate but not univariate information (Coutanche et al. 2013). The latter approach has been criticized for ignoring globally distributed activity patterns and for introducing spatial inaccuracies (Stelzer et al., 2014). Alternatively, whole brain classifiers have been also presented by employing techniques that promote model generalization such as regularization (Churchill et al., 2014; Ryali et al., 2010; Yamashita et al., 2008).

Further concerns regarding importance maps from MVPA classification pertain their extraction, reproducibility and visualization. As univariate feature selection has been criticized for removing multivariate information (Coutanche, 2013), extraction and visualization of importance maps should be also performed in a multivariate manner (Schrouff et al., 2013). Furthermore, since classifiers are typically trained through an optimization process of initially random parameters, multiple runs of the same classifier may generate different importance maps (Rasmussen et al., 2011).

Here we address the aforementioned issues by performing inter-subject whole brain classification of fMRI data. We applied a linear neural network based classifier in three simulated and two different empirical datasets from different domains (emotional states and viewing objects). Subsequently, we extracted importance maps using methods based on classifier weights, weight-input product, output difference and layerwise relevance propagation introduced by Montavon et al. (2017). We applied this scheme in a simulated dataset to demonstrate that importance extraction methods of neural network classifiers can efficiently localize multivariate patterns with high reproducibility. Subsequently, we applied our scheme to two fMRI datasets that have been successfully used for classification. In the first dataset, emotions elicited by short movie clips were classified (Saarimäki et al., 2016). In the second dataset, visual objects were classified during an object recognition task (Haxby et al., 2001). Our results indicate that neural networks succeed in whole-brain classification and identifying involved brain regions with better sensitivity than univariate approaches.

## 2. Methods

### 2.1 Dataset 1: Emotions induced by short movie clips

#### 2.1.1 Participants

Twenty-one volunteers (12 males, ages 19–33, mean age 24.9 years) participated in the experiment. All participants were healthy with normal or corrected-to-normal vision and gave written informed consent.

#### 2.1.2 Design of experiment

For details regarding the experimental protocol, see Saarimäki et al. (2016). Briefly, emotions were induced using short movie clips. Fifty 10-s movie clips were chosen from a video database validated to evoke basic emotions (Tettamanti et al., 2012). We used clips that elicited the most reliable emotions in five emotion categories (10 clips per category): disgust, fear, happiness, sadness, and neutral. The clips were randomly divided into two sets with five movies from each category in both sets. During fMRI, both sets of movie clips were presented twice, thus resulting in four runs in total. Each run lasted for 12min 50s. Each clip was preceded by a 5-s fixation cross and followed by a 15-s washout period. The participants were instructed to view the movies similarly as they would watch TV and to focus on the emotional content of the movie clip. No active task was required during fMRI scanning. The stimuli were delivered using Presentation software (Neurobehavioral Systems Inc., Albany, CA, USA). They were back-projected on a semitransparent screen using a 3-micromirror data projector (Christie X3, Christie Digital Systems Ltd., Mönchengladbach, Germany) and from there via a mirror to the participant. Further details concerning the experiment design and data acquisition can be found in (Saarimäki et al., 2016).

#### 2.1.3 MRI Data Acquisition

MRI data were collected on a 3T Siemens Magnetom Skyra scanner at the Advanced Magnetic Imaging Centre, Aalto NeuroImaging, Aalto University, using a 20-channel Siemens volume coil. Whole-brain functional scans were collected using a whole brain T2^*^-weighted EPI sequence with the following parameters: 33 axial slices, TR = 1.7 s, TE = 24 ms, flip angle = 70°, voxel size = 3.1 × 3.1 × 4.0 mm, matrix size = 64 × 64 × 33, field of view (FOV) = 198.4 mm. A custom-modified bipolar water excitation radio frequency (RF) pulse was used to avoid signal from fat. High-resolution anatomical images with isotropic 1 × 1 × 1 mm voxel size were collected using a T1-weighted MP-RAGE sequence.

### 2.2 Dataset 2: Visual object recognition task

#### 2.2.1 MRI Data Acquisition

This dataset was obtained from the OpenfMRI database (Poldrack & Gorgolewski, 2017; accession number ds000105). Stimuli were gray-scale images of faces, houses, cats, bottles, scissors, shoes, chairs, and nonsense patterns. Control nonsense patterns were phase-scrambled images of the intact objects. Twelve time series were obtained in each subject. Neural responses, as reflected in hemodynamic changes, were measured in six subjects (five female and one male) with gradient echo echo-planar-imaging on a GE 3T scanner (General Electric, Milwaukee, WI) [repetition time (TR) = 2500 ms, 40 3.5-mm-thick sagittal images, FOV = 24 cm, echo time (TE) = 30 ms, flip angle = 90] while they performed a one-back repetition detection task. High-resolution T1-weighted spoiled gradient recall (SPGR) images were obtained for each subject to provide detailed anatomy (124 1.2-mm-thick sagittal images, FOV = 24 cm). Further details regarding its acquisition can be found in https://openfmri.org/dataset/ds000105/ as well as in the original publication (Haxby et al., 2001). Since the 12th run was missing from subject 5 in the open dataset, the 12th run was excluded from all subjects to achieve equal number of samples per subject.

### 2.3 Data preprocessing

Data were preprocessed using FSL 5.0 (Jenkinson et al., 2012; Smith et al., 2004; Woolrich et al., 2009). Motion was corrected using MCFLIRT (Jenkinson and Smith, 2001; Jenkinson et al., 2002) and non-brain matter was removed using BET (Smith, 2002). High-pass temporal filtering was applied using Gaussian-weighted least-squares straight line fitting with sigma of 55 volumes. For inter-subject classification, the functional data were registered to 2 × 2 × 2 mm MNI152 standard space template using FLIRT (Jenkinson and Smith, 2001; Jenkinson et al., 2002). The brain-extracted T1-weighted images were first normalized to the MNI space and the normalization parameters were subsequently applied to the EPI images. All registrations were performed using 9 degrees of freedom. No spatial smoothing was applied.

Framewise displacement (FD), was calculated for each subject as suggested by Power et al. (2012). All subjects in both datasets had more than 90% of time points with framewise displacement (FD) less than 0.5 mm. Average FD was 0.12mm and 0.07mm for dataset 1 and dataset 2 respectively.

In both datasets, a 2 × 2 × 2 mm MNI152 standard brain mask was used. To reduce the number of voxels in the analysis, we performed spatial downsampling to 4 × 4 × 4 mm voxels to the EPI data as well as the binary mask. This resulted to a total number of 28 586 voxels.

Average activation maps were used as input to the classifier. Specifically, for dataset 1, we used the temporal average over an 11.9 second interval (7 TRs) centered around the end of each movie clip (emotional peak experience). For dataset 2, we used a 12.5 second interval (5 TRs) from stimulus onset.

### 2.4 Data preparation for simulations

The short movie clips dataset was used as the basis for the simulated data. For each time point we performed random permutations of the voxels for each sample. To ensure there are no consistent mean effects (Junghöfer et al., 2015; Hayasaka 2013), all the samples were also randomly reordered between categories. The result was used as a basis for generating different simulation scenarios.

### 2.5 Simulation scenarios

We generated patterns of different spatial size and amounts of overlap. More specifically we simulated partly overlapping patterns, completely overlapping patterns, large patterns (one 5th of the total number of voxels), small patterns (one 100th of the total number of voxels) and patterns where the voxels are a subset of some other category’s pattern. Univariate effects were generated by adding normally distributed noise with a low mean to avoid high classification performance. Three simulation scenarios were implemented with increasing complexity. The three simulation scenarios are presented in Figure 1c. More specifically the three scenarios were designed as following:

**Figure 1:**
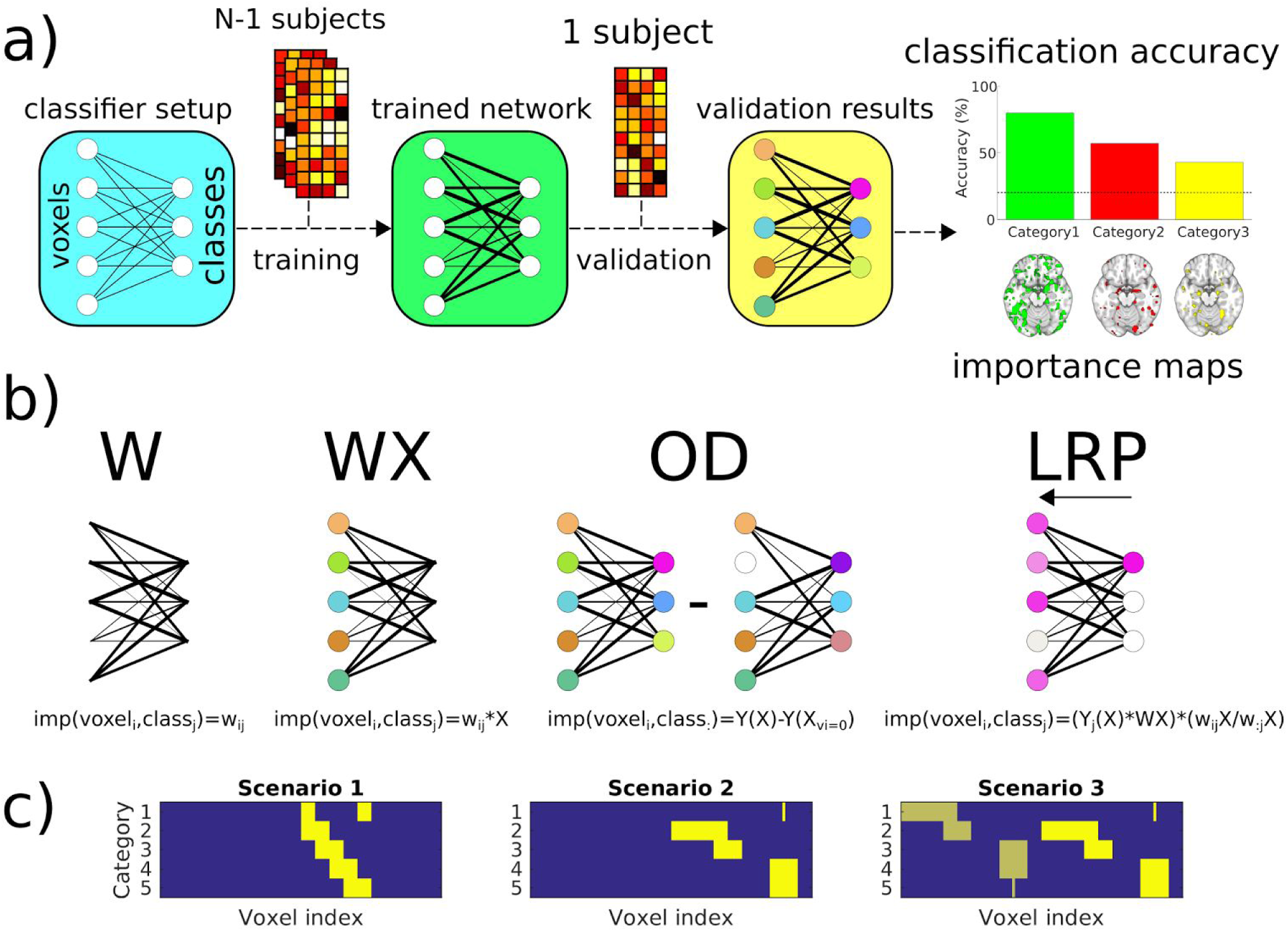
Classification analysis workflow (a), visual representation of the four importance extraction methods (b) and ground truth for each simulation scenario (c).

**Scenario 1**: Overlapping patterns with same size: Each category consisted of patterns with the size of one tenth of the total number of voxels. Each category had 50% overlap with the next category and the other 50% with the previous category. No voxels were important for only one category.

**Scenario 2**: Complex patterns - same magnitudes: Different effect sizes were chosen for each category. The effect size for category 1 was one hundredth of the total number of voxels. Category 2 consisted of one fifth of the total number of voxels. Categories 3, 4 and 5 consisted of patterns with size one tenth of the total number of voxels. Category 3 had 50% overlap with category 2. Categories 4 and 5 were fully overlapping, while the pattern for category 1 was a subset of the voxels of category 4 and 5. Gaussian noise with mean 0.05 and unit variance was added to the regions.

**Scenario 3**: Complex patterns - multiple magnitudes: This scenario incorporates the patterns of Scenario 2 two times the Scenario 2 with different magnitudes. The patterns are also shifted among categories so that there are different effect sizes for each category. Gaussian noise with mean 0.05 and unit variance were added on the left side. On the right side the mean was 0.07 (see Scenario 3 in Figure 1c).

### 2.6 Univariate tests for activation differences - Student’s T-test

We applied unpaired two sample t-tests to examine univariate activation differences. For each voxel, we performed one-versus-rest comparisons, that is contrasting all the samples of one category versus the samples of the rest categories. We extracted p-values as well as t-values for each voxel and each category.

### 2.7 Classifier setup

Artificial neural network based classifiers were used for the classification as implemented in a neural network toolbox for developed by Lapuschkin et al. (2016). The classifier had no hidden layers. The classifier utilized softmax activation function in the output layer. A low minibatch size of 20 was selected in order to avoid overfitting (Keskar et al., 2017). Training was performed for 10000 epochs using stochastic gradient descent as an optimization algorithm. Learning factor was set by default to 0.005. The L1 norm was used as a loss function, which has shown shown increased robustness compared to other loss functions in neural networks (Gorban et al., 2016; Wang et al., 2008; Zhao et al., 2015). The model was trained using backpropagation and stochastic gradient descent (SGD) was selected as the optimization algorithm. (LeCun et al., 2012). The process was repeated for 1000 times. Leave-one-subject-out (LOSO) cross-validation was used for the evaluation of the trained classifier, both in terms of classification accuracy as well as for extracting importance maps. More specifically, the data were split to a training set consisting of all the samples from all but one subjects and a validation set consisting of the samples from the left-out subject. This process was repeated so that all subjects were used as left-out subjects. MATLAB (MATLAB 2016b, The MathWorks, Inc., Natick, Massachusetts, United States) was used for the classification as well as for all steps of data analysis and visualization.

### 2.8 Extracting voxel importances

Four methods were tested for importance extraction. The first one uses only the weights of the trained classifier in a similar fashion as suggested in Pereira et al. (2009) and here is denoted as **W**. The second one relies on the weights-activations product as proposed by Polyn et al. (2005), here denoted as **WX**. The third method measures the difference in the output of the classifier after removing one voxel. This process was repeated for each voxel. Since this measure measures the classifier’s output difference, we refer to it as **OD**. As the output of the classifier ranges from 0 to 1, importances extracted by the OD method range from −1 to +1, reflecting the two extreme cases, where classification depends only on one input and upon its removal the output changes from 0 to 1 or from 1 to 0 respectively. In practice the values are much lower as their magnitude depends on the output as well as on the number of variables that contribute to the classifier. The fourth importance extraction method decomposes the classifier’s output to the inputs through layerwise relevance propagation (**LRP**) as introduced by (Bach et al., 2015). See Figure 1b for a visual representation of the 4 methods.

### 2.9 Statistical evaluation of classification results

#### 2.9.1 Permutation runs and significance threshold

To generate a null-distribution for classification accuracies as well as for importance maps, we performed permutations by running the classifier after shuffling the output labels. More specifically, the labels were shuffled before splitting data to training and validation sets, therefore all labels were shuffled. This process was performed for 1000 times. The resulting permutations were used for contrasting classification accuracies against a null-distribution, as well as for setting a significance threshold for the importance maps.

#### 2.9.2 Measuring pattern reproducibility

After extracting a significance threshold for classification accuracy and importance maps, a reproducibility measure was defined by measuring the number of times a voxel appeared significant within the 1000 runs of the classifier. This resulted to reproducibility curves showing the number of voxels that appeared significant for a certain number of runs.

#### 2.9.3 Reproducibility permutations and reproducibility threshold

To test for false positives, that is, the number of voxels that appear significantly reproducible by chance, a second permutation set was generated in an identical manner as in significance threshold permutations. Since no voxels are expected to exceed significance threshold in permutations, the reproducibility threshold was selected so that no voxels from the reproducibility permutations appeared significant.

## 3. Results

### 3.1 Classification accuracies & confusion matrices

Average classification accuracy, classification accuracy per category and confusion matrices are summarized in Figure 2 for each scenario and each dataset.

**Figure 2.**
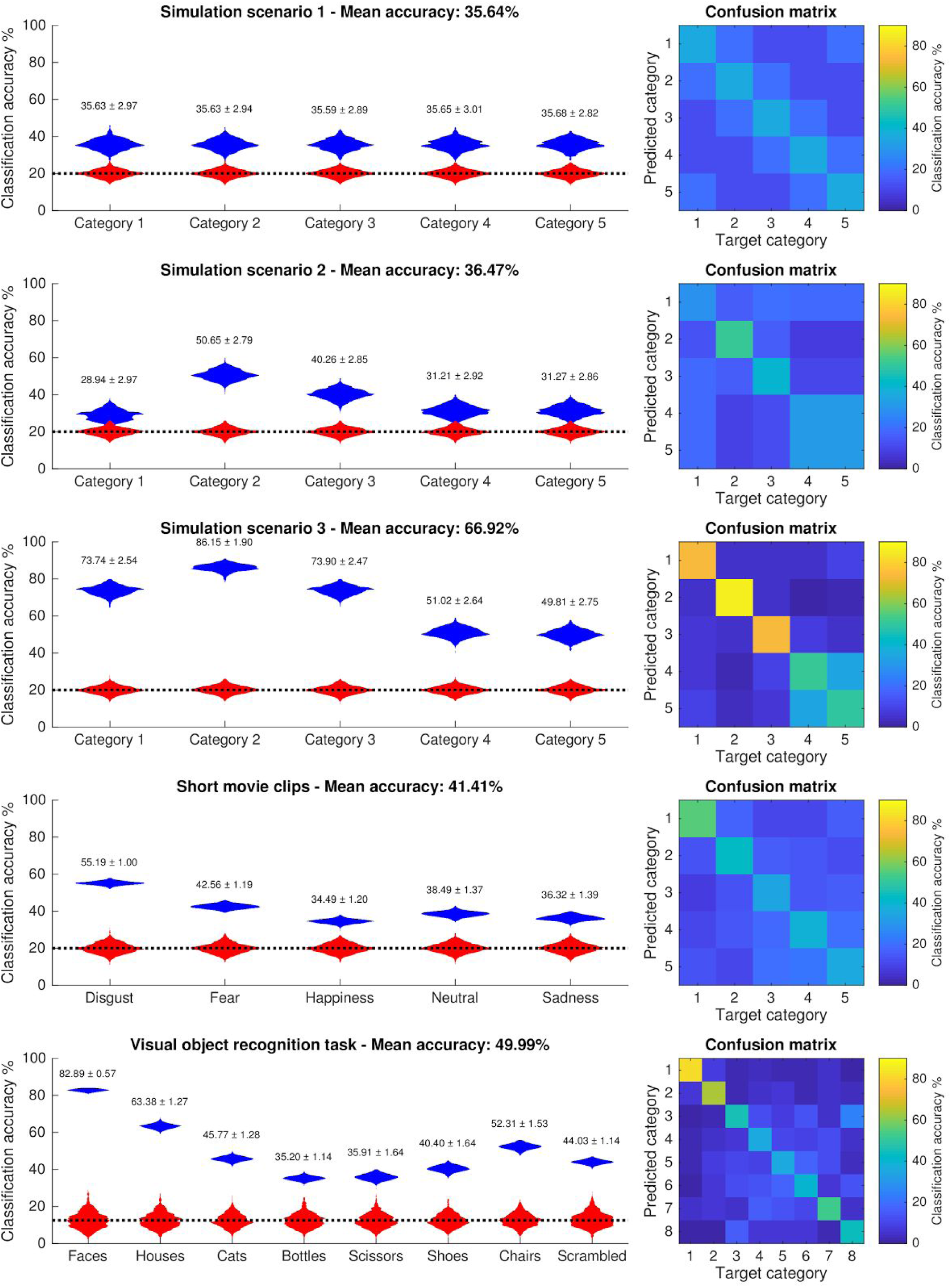
Classification accuracy violin plots and confusion matrix for each simulation scenario and each dataset. Classification accuracy distributions are shown in blue and red for classification runs and permutation runs respectively. The horizontal dashed line corresponds to the theoretical chancel level accuracy. Numbers above each distribution denote category-specific mean and standard deviation of classification accuracy. All distribution differences are statistically significant at p<0.001.

### 3.2 T-test results

Univariate activation differences were calculated by applying a two sample t-test in a one-versus-all fashion. T-values were obtained for each voxel and each category. Results are shown in Figure 3. Although t-values visually indicate the important regions, the values are not high enough to survive any sensible statistical threshold.

**Figure 3:**
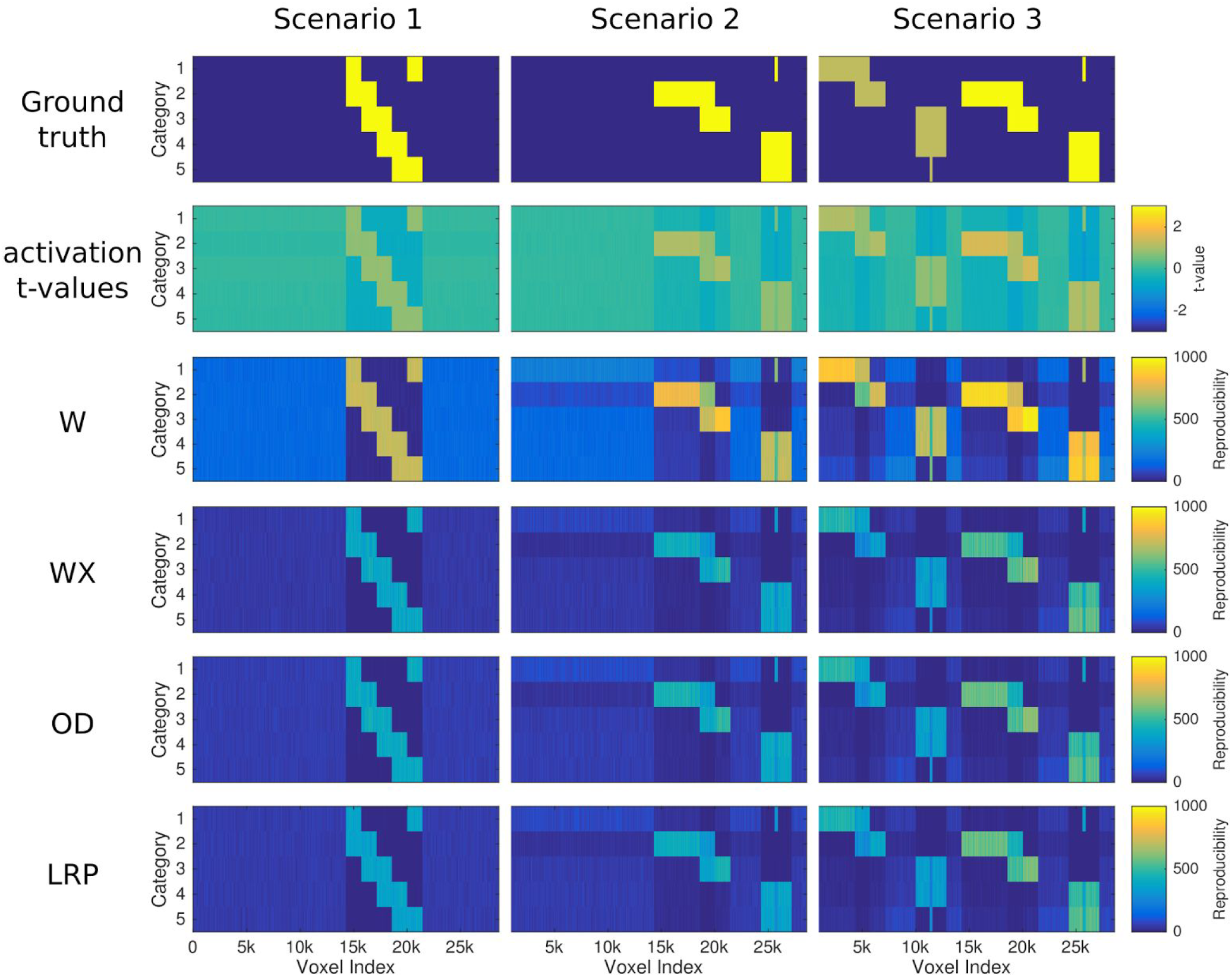
Activation t-test and importance reproducibility results for each simulation scenario. First row shows the ground truth for each scenario. Second row shows activation t-values when contrasting one-versus-all categories. The last four rows show reproducibility values for each importance extraction method, i.e. how many times each voxel appeared significant (p<0.01) out of 1000 runs. Univariate t-values are low and do not survive any significance threshold, although visually they indicate important regions.

### 3.3 Reproducibility maps, statistical maps & brain maps

Reproducibility maps were generated by enumerating for each voxel the number of times it appeared significant out of the 1000 runs of the classifier. Reproducibility maps for the simulation scenarios are shown in Figure 3. An example of importance reproducibility maps for each empirical dataset is shown in Figure 4; the number of overlapping methods is presented for a reproducibility threshold of 500 and significance threshold p<0.01. This analysis workflow generated numerous brain maps for each dataset, each importance extraction and each category. T-value maps were generated from the univariate activation t-tests. Importance maps, averaged over the 1000 runs were also generated as well as reproducibility maps for p<0.01. For better visualization and inspection of the results for the two empirical datasets, two brain map collections were created in NeuroVault, one for the short movie clips dataset (https://neurovault.org/collections/3032/) and one for the visual object recognition dataset (https://neurovault.org/collections/3033/). Since the actual importance values can be too low to be properly shown in NIFTI format, all importance values were multiplied by 1000. All brain maps are in 4×4×4 mm resolution.

**Figure 4.**
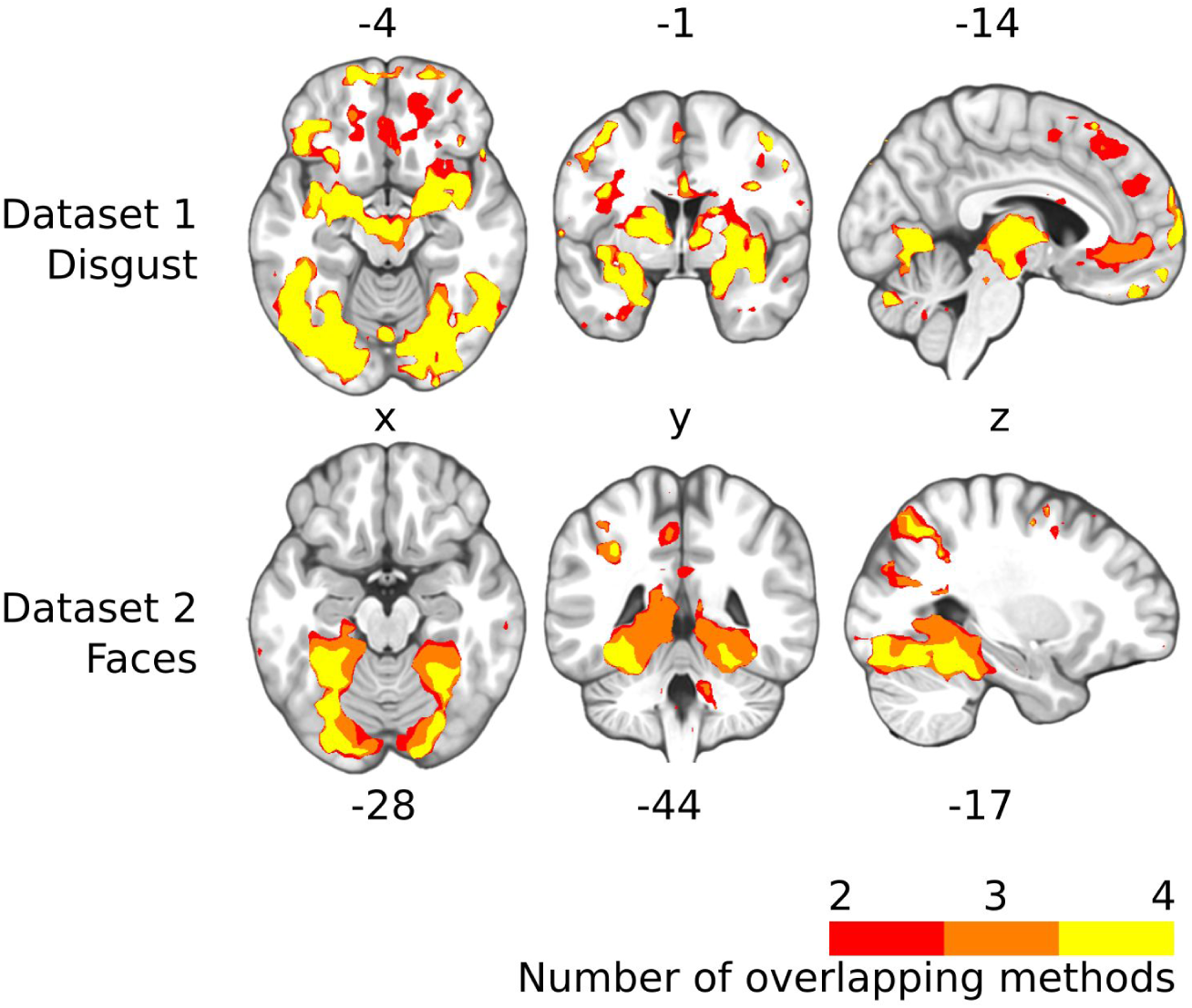
Example importance maps for each empirical dataset. The category with the highest classification accuracy is presented for each dataset (“disgust” for dataset 1 and “faces” for dataset 2). Each map represents the number of methods that exceed reproducibility threshold of 500 in significance threshold at p<0.01. The slices follow the neurological convention (right is right) and locations are shown in MNI coordinates.

### 3.4 Reproducibility curves

For each importance extraction method reproducibility curves were generated indicating the number of voxels that exceed the significance threshold generated by permutations. Reproducibility curves were also generated for the reproducibility permutations. Reproducibility curves for each simulation scenario and each dataset are presented in Figure 5. For the three simulation scenarios where ground truth is known, voxel accuracy was measured (i.e. the percentage of correctly defined voxels as important or not) as well as False Discovery Rate (FDR) and False Omission Rate (FOR). The results for different significance thresholds are shown in Supplementary Figure 2.

**Figure 5:**
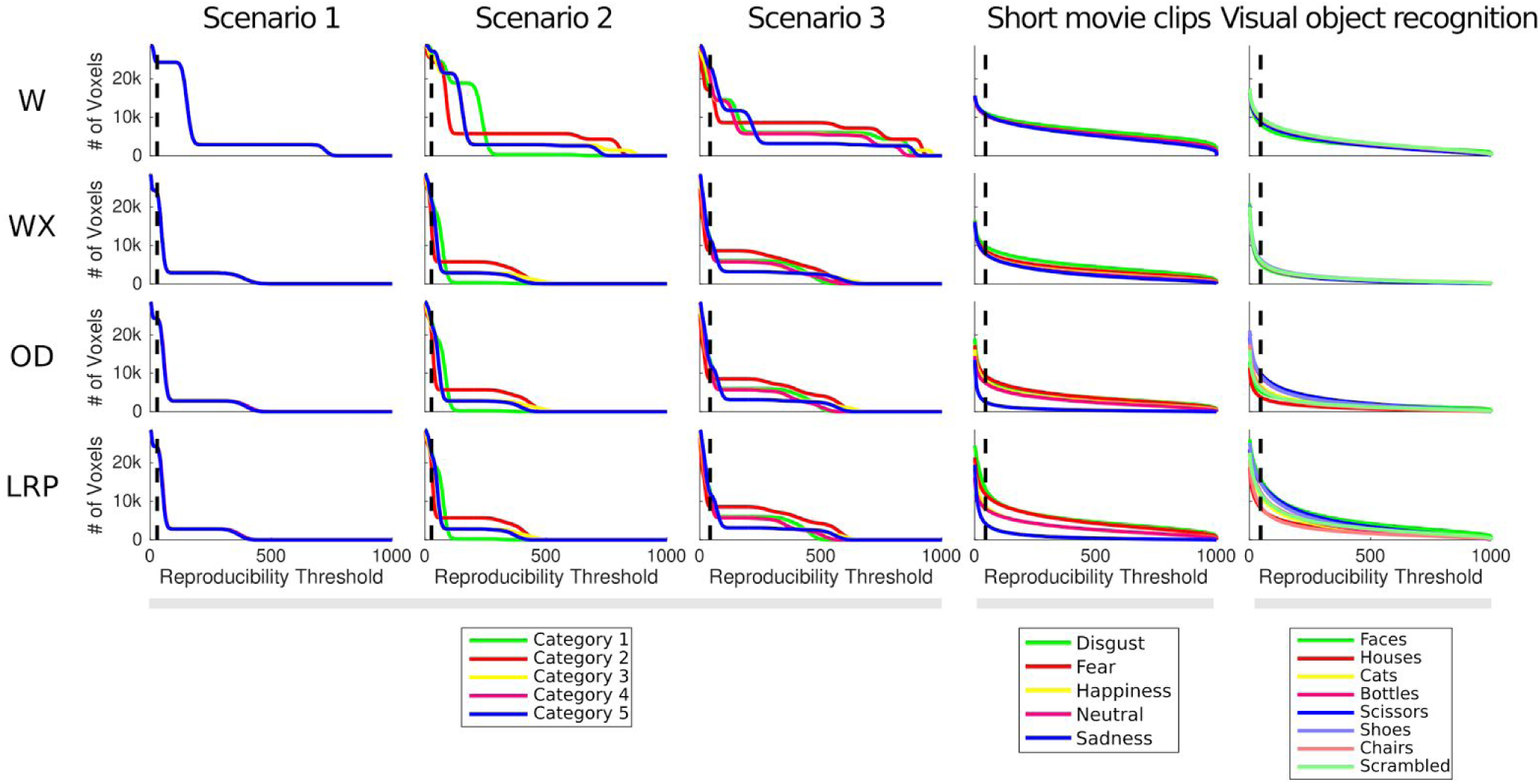
Reproducibility curves for extracted importances for all simulation scenarios and datasets (p<0.01). Vertical dashed lines indicate the reproducibility threshold where zero voxels appear significant in the reproducibility permutations. The reproducibility threshold generated by permutations is too lenient, especially for the W method.

### 3.5 Univariate versus multivariate information

Reproducibility plots versus univariate t-values plots were generated to examine the relation between univariate and multivariate information. Figure 6 depicts a representative example of low univariate information (absolute t-value<1, degrees of freedom = 2098, p<0.15) and high reproducibility (>500). The activation t-value versus importance reproducibility plots can be found in the Supplementary Figure 1.

**Figure 6.**
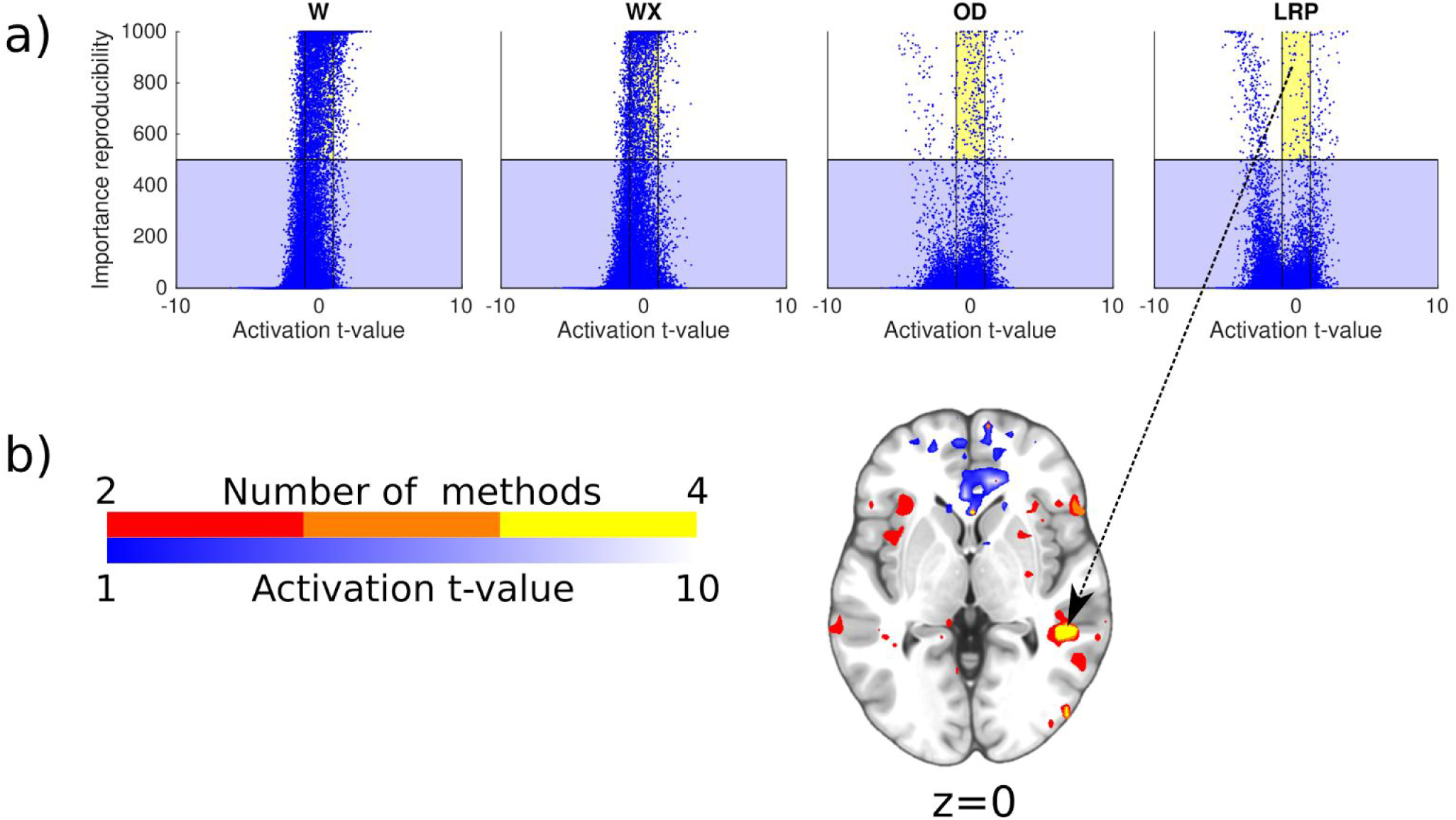
Example of low univariate information but high importance reproducibility. (a) Univariate t-values versus importance reproducibility plots for category “sadness” in the short movie clips dataset. Each dot represents a voxel. Yellow area indicates reproducibility > 500 (with p<0.01) and low univariate t-value (<1; 2098 degrees of freedom), (b) A region in the right superior temporal sulcus that exhibits highly reproducibility but low univariate information for all importance extraction methods. Absolute t-values higher than 1 are shown in blue-white gradient.The number of methods that exceed a reproducibility threshold of 500 are shown in red-yellow discrete gradient.The slice follows the neurological convention (right is right).

## 4. Discussion

In this paper, we provided a better understanding of neural network based fMRI classification using importance extraction methods. The methods were validated using simulation scenarios to examine their behaviour in terms of importance reproducibility. The resulting reproducibility maps for the two datasets we examined show high similarity to univariate statistics but with increased statistical power. A particularly interesting case is the combination of high reproducibility but low univariate values (see Figure 6), which indicates complex interaction of voxel activations. Interpreting such interactions requires further research but it is important to underline that such voxels could be excluded if a univariate feature selection method was applied prior to classification.

### 4.1 Classifier selection

Neural network classifiers have been previously used to classify fMRI data either with hidden layers (Bertolino et al., 2014; Floren et al., 2015; Misaki et al., 2006) or without (Polyn et al., 2005; Saarimäki et al., 2016). The majority of MVPA studies use support vector classifiers (SVC) (Cox & Savoy, 2003; De Martino et al., 2008, Ethofer et al., 2009; Habes et al., 2013; LaConte et al., 2005; Kamitani & Tong, 2005; Lahnakoski et al., 2014; Lie et al., 2013; Meier et al., 2012; Mourão-Miranda et al., 2005; Mourão-Miranda et al., 2007; Rasmussen et al., 2011; see also Sundermann et al., 2014, for an extended list) due to fast training and good performance in ill-posed problems such as in fMRI classification (Etzel et al., 2013). Main drawback is that SVCs are inherently binary classifiers, hence not optimal for multiclass problems. There are variations that face this limitation, typically either by performing classifications between each category pair or by one-versus-all (OVA) classification (Bishop, 2006). Although evaluation of similar importance extraction approaches for other classifiers would allow a more general evaluation of methods and provide further insight, this would require reformulation of the analysis that would hinder the interpretability of the results. Since methods for neural networks are under intense development due to their impressive performance in several fields, we predict that there will be an increase of their application in neuroscience.

### 4.2 Classification accuracy and confusion matrices

#### 4.2.1 Simulation scenario 1

In the first simulation scenario the effect size and the effect magnitude for all categories were identical. Therefore we expected a similar classification accuracy for all categories. Since each category shares overlapping representation with two other categories, misclassifications were expected to be prominent. Although the effect was non-linear – since coactivation of two regions is required for each category – the linear classifier managed to detect this effect to some extent. This ability of linear classifiers to partly detect non-linear relationships has been addressed also by Davis et al. (2014). Our results demonstrate that linear classifiers can indeed detect effects that rely on mutual activation of two or more regions. We however expect that non-linear classifiers would show higher flexibility in the decision boundary and hence better performance, yet this remains to be tested in future studies.

#### 4.2.2 Simulation scenario 2

Classification accuracies in the second scenario show that performance depends on the effect size, which is also the main benefit expected from MVPA. When the effect magnitude is identical, classification accuracy is proportional to the effect size. Another observation is that categories 4 and 5 exhibit significant classification accuracy although they consist of identical patterns. This reveals the need to examine confusion matrices, since the two classes are misclassified among each other but are well discriminated from the rest, leading to significant classification accuracy (see Figure 2).

#### 4.2.3 Simulation scenario 3

This scenario was most similar to real data, since it incorporates both univariate and multivariate effects, different effect sizes, as well as different magnitudes. Overall, the classification performance was best of all the simulated scenarios since, this scenario contains similar information as scenario two plus more patterns with higher magnitude. Category 2 exhibits the highest classification accuracy. Although categories 3 and 4 have the same effect size, category 3 shows higher classification accuracy, attributed to the voxels that are active exclusively for that category (see pattern of category 3 in Figure 1c). The results indicate that classification accuracy is proportional both to effect magnitude and effect size but inferring which is the case is not trivial.

### 4.3 Interpretability of importance maps

Although the weights of the classifier (presented as “W” in Figure 1b) may constitute the most intuitive approach to estimate importances of a linear classifier, there are a number of disadvantages. First, there is no direct interpretation of the magnitude and sign of the weights. For example, a negative sign indicates that increasing the voxel activity causes a decrease in the classifier output. Thus, the contribution of a given voxel, whether it is positive or negative, depends on the sign of the activity (e.g. negative input and negative weight contribute positively to the output). While from neuroscientific perspective, of course, both “activations” and “deactivations” do carry meaningful information (given that both constitute modulations of spontaneous activity and are thus constituents of functional brain states), being able to distinguish between these two would be desirable. The second and more important disadvantage comes from that the weights are defined during training and hence are prone to overfitting. The WX method solves the sign interpretability issue, as well as the latter problem, since the validation set is used as input. However, there is no quantitative interpretation of the importances in the WX method. The OD method solves the problem of quantitative interpretability since the OD importances range from −1 to 1, indicating the change of the classifier output when a certain voxel is removed. Furthermore, it can be easily implemented and tested in other classifiers. However, all the previous methods have two disadvantages. First, interpretation of a multivariate classifier is derived in a univariate manner; each voxel importance is estimated separately ignoring of the rest of the voxels. This issue has also been mentioned in a previous study (Schrouff et al., 2013). Second, they do not take into account the actual output of the classifier, that is, how well the validation set was classified. These two issues are addressed by the LRP approach as the classifier’s output is distributed back to the inputs. Since the output is redistributed to the inputs, the sum of the importances equals the output of the classifier, providing a direct interpretation of each importance map.

### 4.4 Thresholding of importance maps

Thresholding and visualizing importance maps has been a common practice in MVPA studies (McDuff et al., 2009; Rissmann et al., 2010; Saarimäki et al., 2016) although it has been criticized as inappropriate since thresholding multivariate information in a univariate manner is a questionable practice (Schrouff et al., 2013). While being aware of this potential pitfall, thresholding of multivariate maps serves two major functions. First, thresholds generated by permutations indicate a value that is statistically unlikely to be result of a random classifier. Second, thresholding provides easier visualization of importance maps. Another important issue that to our knowledge has not been discussed in MVPA community is whether importance maps resemble the activation patterns per se or rather indicate the localization of the patterns but not the activation patterns per se. In the latter case, thresholding is a rational approach to follow. Previous work on image classification, using layerwise relevance propagation, has shown that importance maps indicate important features regardless of the intensity of the input (Bach et al., 2015; Montavon et al., 2017).

### 4.5 Generating significance thresholds

Permutation testing is an established approach for significance testing due to its intuitive and non-parametric approach while minimizing assumptions of the model (Stelzer et al., 2013). Its main drawback is its computational complexity. Furthermore, since type I errors have emerged as a major pitfall in fMRI analysis (Eklund et al., 2016; Lieberman & Cunningham, 2009), larger scale analyses, where thousands of voxels are involved, require a proportionally higher number of permutations to test for multiple comparisons. This may require an intractable amount of computations. We performed 1000 permutations to extract significance thresholds for the importances of the 28586 voxels. The expected number of false positives at a significance level of p=0.01 is ∼280 voxels. To minimize the number of false positive results, we introduced a set of reproducibility permutations where the number of false positive occurrences is measured per voxels. Reproducibility of importance maps has been addressed earlier by Rasmussen et al. (2011), showing that under certain circumstances different classification runs may yield similar classification accuracies but different importance maps.

### 4.6 Limitations

#### 4.6.1 Time point selection

There was no performance-driven motivation in the selection of time points for the analysis of both datasets. Even if time point selection is not optimal, the interpretation of the results is independent of the selected time points and is not expected to bias towards any direction.

#### 4.6.2 Simulated dataset and limitations

Although our simulation datasets were generated to resemble as well as possible real fMRI data there are certain differences and limitations. More specifically, our simulations did not address spatial differences between subjects. Furthermore, signal quality differences that exist between regions of the brain (e.g. SNR of cortical and subcortical regions) were not taken into account. Although different effect magnitudes were simulated (see simulation scenario 3, Figure 1c), real datasets are expected to consist of a wider and continuous range of effect magnitudes. Furthermore, there is no proof that activations follow a gaussian distribution like the effects generated in our simulations. However, the non-parametric nature of the statistical methods we used does not introduce any distribution related bias. Being aware of these existing limitations, conclusions regarding the statistical power of importance maps in comparison to univariate statistics can be still safely drawn.

#### 4.6.2 Inter-subject versus within-subject classification

Intersubject classification has shown low performance compared to within-subject due to variability in subjective experiences, spatial inaccuracies introduced by anatomical differences and inaccuracies due to registration to a brain template (Haxby et al., 2014). There have been a few approaches suggested to tackle such this problem, either through coregistration based on functional connectivity (Conroy et al., 2013) or through hyperalignment (Haxby et al., 2011). Being aware of these inaccuracies, in this study we focus on inter-subject classification for two major reasons. First, we reckon inter-subject classification of high significance both in research and clinical setup, as it addresses beyond subject-specific commonalities, given the existing limitations. Second, the proposed setup exploits the full dataset, leading to more samples per input, which is a desirable feature while training classification models. However, the classification analysis workflow for intra-subject classification would be identical, requiring only different segmentation of the dataset (e.g. leave-one-run-out setup). Hence, the applied LOSO cross-validation tested whether the decoded patterns generalized across subjects.

## 5. Conclusions

The increasing use of classification tools in fMRI data analysis has necessitated methods that interpret the classifiers' decisions with regard to the classifier input. Such methods are in the spotlight of machine learning research and we showed that they are directly applicable to fMRI classification. Our findings demonstrate the increased statistical sensitivity of such methods compared to univariate approaches and provide a better understanding of the classifiers' behaviour in the form of importance maps. Brain regions that exhibit high importance but low univariate information are of particular interest and require further research to interpret the underlying mechanisms from a neuroscientific perspective.

## Acknowledgements

We thank Marita Kattelus for her help with the data acquisition. We also acknowledge the computational resources provided by the Aalto Science-IT project.

## Supplementary material

**Supplementary figure 1:**
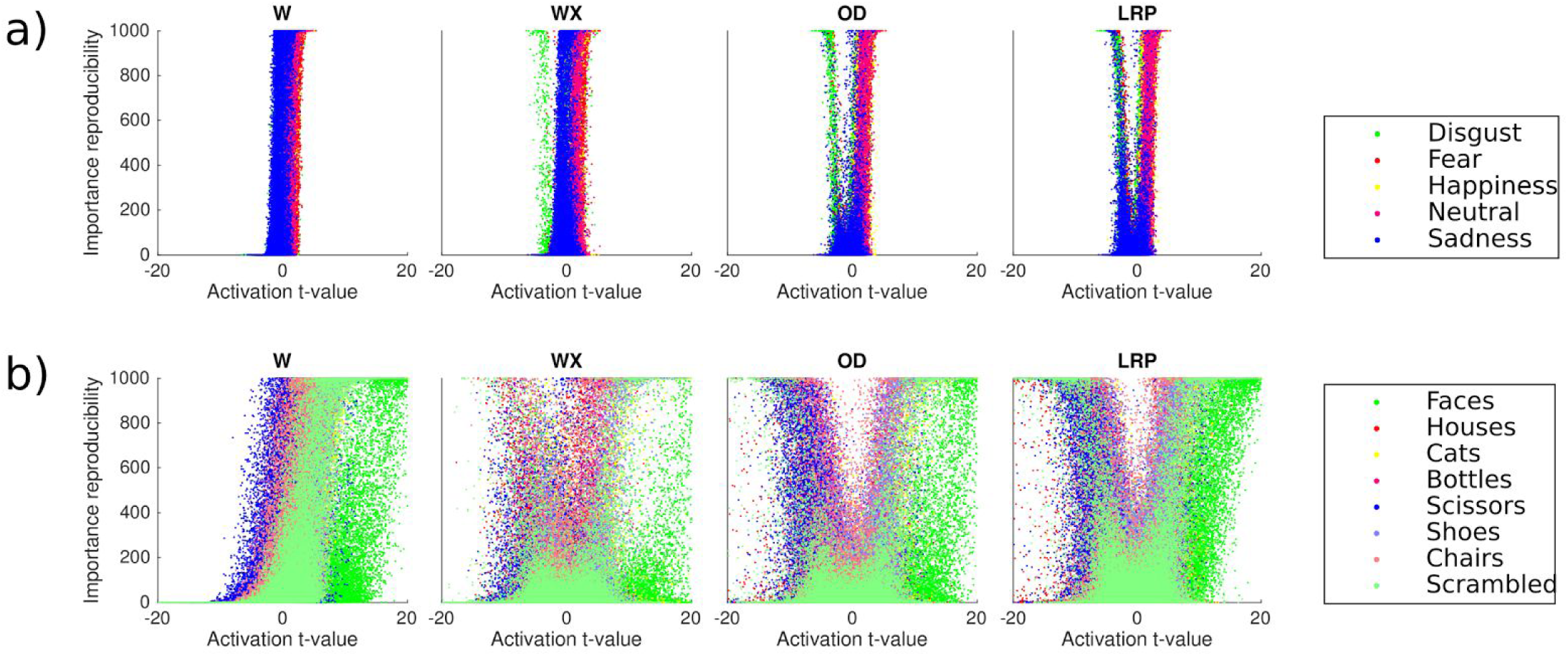
Activation t-value versus importance reproducibility plots. Short movie clips dataset (a) and visual object recognition task (b) for p<0.01. The categories with the highest classification accuracy in each dataset exhibit the largest t-values hinting towards high discriminability under the presence of strong univariate effects. For OD and LRP methods, more negative t-values are assigned high importance reproducibility values.

**Supplementary figure 2:**
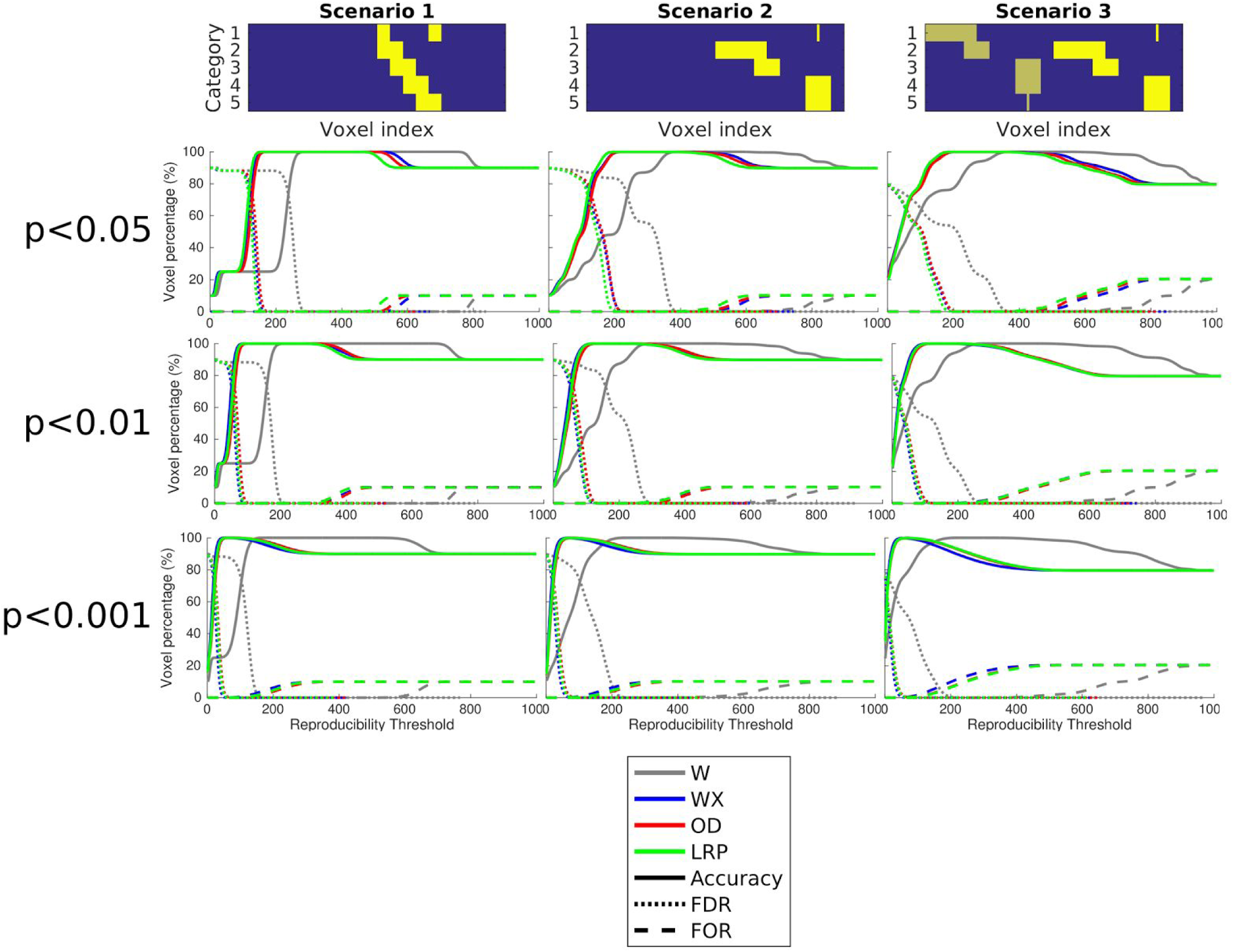
Voxel accuracy, false discovery rate (FDR) and false omission rate (FOR). For each significance threshold (p=0.001, 0.01, 0.05), each simulation scenario and each importance extraction method. Methods WX, OD and LRP exhibit similar behavior in terms of voxel accuracy, FDR and FOR, while W method is more lenient and thus requires higher reproducibility threshold. The reproducibility curve and reproducibility threshold for W is more dependent on the dataset (reproducibility threshold around 300 for scenario 1 and 400 for scenario 3 when p<0.05) compared to the rest of methods.

## References

Bach, S., Binder, A., Montavon, G., Klauschen, F., Müller, K.R., Samek, W., 2015. On pixel-wise explanations for non-linear classifier decisions by layer-wise relevance propagation. PLoS One 10, 1–46.

Bertolino, N., Ferraro, S., Nigri, A., Bruzzone, M.G., Ghielmetti, F., 2014. A neural network approach to fMRI binocular visual rivalry task analysis. PLoS One 9.

Bishop, C. M. 2006. Pattern Recognition and Machine Learning. Springer. pp. 182–183.

Churchill, N.W., Yourganov, G., Strother, S.C., 2014. Comparing within-subject classification and regularization methods in fMRI for large and small sample sizes. Hum. Brain Mapp. 35, 4499–4517.

Cohen, J.D., Daw, N., Engelhardt, B., Hasson, U., Li, K., Niv, Y., Norman, K.A., Pillow, J., Ramadge, P.J., Turk-Browne, N.B., Willke, T.L., 2017. Computational approaches to fMRI analysis. Nat. Neurosci. 20, 304–313.

Conroy, B.R., Singer, B.D., Guntupalli, J.S., Ramadge, P.J., Haxby, J. V, 2013. Inter-subject alignment of human cortical anatomy using functional connectivity. Neuroimage 81, 400–411.

Coutanche, M.N., 2013. Distinguishing multi-voxel patterns and mean activation: why, how, and what does it tell us? Cogn. Affect. Behav. Neurosci. 13, 667–73.doi:10.3758/s13415-013-0186-2

Coutanche, M.N., Thompson-Schill, S.L., Schultz, R.T., 2011. Multi-voxel pattern analysis of fMRI data predicts clinical symptom severity. Neuroimage 57, 113–23.

Cox, D.D., Savoy, R.L., 2003. Functional magnetic resonance imaging (fMRI) “brain reading”: detecting and classifying distributed patterns of fMRI activity in human visual cortex. Neuroimage 19, 261–270.

Davis, T., LaRocque, K.F., Mumford, J.A., Norman, K.A., Wagner, A.D., Poldrack, R.A. (2014). What do differences between multi-voxel and univariate analysis mean? How subject-, voxel-, and trial-level variance impact fMRI analysis. NeuroImage, 97, 271–283. https://doi.org/10.1016/j.neuroimage.2014.04.037

De Martino, F., Valente, G., Staeren, N., Ashburner, J., Goebel, R., Formisano, E., 2008. Combining multivariate voxel selection and support vector machines for mapping and classification of fMRI spatial patterns. Neuroimage 43, 44–58. doi:10.1016/j.neuroimage.2008.06.037

Eklund, A., Nichols, T.E., Knutsson, H., 2016. Cluster failure: Why fMRI inferences for spatial extent have inflated false-positive rates, Proceedings of the National Academy of Sciences. doi:10.1073/pnas.1602413113

Ethofer, T., Van De Ville, D., Scherer, K., Vuilleumier, P., 2009. Decoding of emotional information in voice-sensitive cortices. Curr. Biol. 19, 1028–33.doi:10.1016/j.cub.2009.04.054

Etzel, J.A., Zacks, J.M., Braver, T.S., 2013. Searchlight analysis: Promise, pitfalls, and potential. Neuroimage 78, 261–269. doi:10.1016/j.neuroimage.2013.03.041

Floren, A., Naylor, B., Miikkulainen, R., Ress, D., 2015. Accurately decoding visual information from fMRI data obtained in a realistic virtual environment. Front. Hum. Neurosci. 9, 327. doi:10.3389/fnhum.2015.00327

Gorban, A.N., Mirkes, E.M., Zinovyev, A., 2016. Piece-wise quadratic approximations of arbitrary error functions for fast and robust machine learning. Neural Networks 84, 28–38. doi:10.1016/j.neunet.2016.08.007

Habes, I., Krall, S.C., Johnston, S.J., Yuen, K.S.L., Healy, D., Goebel, R., Sorger, B., Linden, D.E.J., 2013. Pattern classification of valence in depression. NeuroImage. Clin. 2, 675–83. doi:10.1016/j.nicl.2013.05.001

Haxby, J. V, Gobbini, M.I., Furey, M.L., Ishai, a, Schouten, J.L., Pietrini, P., 2001. Distributed and overlapping representations of faces and objects in ventral temporal cortex. Science 293, 2425–30.doi:10.1126/science.1063736

Haxby, J. V, Guntupalli, J.S., Connolly, A.C., Halchenko, Y.O., Conroy, B.R., Gobbini, M.I., Hanke, M., Ramadge, P.J., 2011. A common, high-dimensional model of the representational space in human ventral temporal cortex. Neuron 72, 404–416.

Haxby, J. V, Connolly, A.C., Guntupalli, J.S., 2014. Decoding Neural Representational Spaces Using Multivariate Pattern Analysis. Annu. Rev. Neurosci. 435–456.

Hayasaka, S., 2013. Functional connectivity networks with and without global signal correction. Front. Hum. Neurosci. 7, 880.

Jenkinson, M., Smith, S.M., 2001. A global optimisation method for robust affine registration of brain images. Med. Image Anal., 5, 143–56.

Jenkinson, Mark, Bannister, P. R., Brady, J. M., Smith, S.M., 2002. Improved Optimization for the Robust and Accurate Linear Registration and Motion Correction of Brain Images. Neuroimage, 17, 825–841.

Jenkinson, Mark, Beckmann, C.F., Behrens, T.E.J., Woolrich, M.W., and Smith, S.M., 2012. FSL. Neuroimage, 62, 782–90.

Junghöfer, M., Schupp, H.T., Stark, R., Vaitl, D., 2005. Neuroimaging of emotion: Empirical effects of proportional global signal scaling in fMRI data analysis. Neuroimage 25, 520–526.

Kamitani, Y., Tong, F., 2005. Decoding the visual and subjective contents of the human brain. Nat. Neurosci. 8, 679–685.

Keskar, N. S., Mudigere, D., Nocedal, J., Smelyanskiy, M., Tang, P. T. P., 2017. On large-batch training for deep learning: Generalization gap and sharp minima. In International Conference on Learning Representations (ICLR)

Kober, H., Barrett, L.F., Joseph, J., Bliss-Moreau, E., Lindquist, K., Wager, T.D., 2008. Functional grouping and cortical-subcortical interactions in emotion: a meta-analysis of neuroimaging studies. Neuroimage 42, 998–1031.

Kohler, P.J., Fogelson, S. V, Reavis, E. a, Meng, M., Guntupalli, J.S., Hanke, M., Halchenko, Y.O., Connolly, A.C., Haxby, J. V, Tse, P.U., 2013. Pattern classification precedes region-average hemodynamic response in early visual cortex. Neuroimage 78, 249–60.

LaConte, S., Strother, S., Cherkassky, V., Anderson, J., Hu, X., 2005. Support vector machines for temporal classification of block design fMRI data. Neuroimage 26, 317–329.

Lahnakoski, J.M., Glerean, E., Jääskeläinen, I.P., Hyönä, J., Hari, R., Sams, M., Nummenmaa, L., 2014. Synchronous brain activity across individuals underlies shared psychological perspectives. Neuroimage 100, 316–324.

Lapuschkin, S., Binder, A., 2016. The LRP Toolbox for Artificial Neural Networks 17, 1–5.

LeCun, Y., Bottou, L., Orr, G. B., Müller K. R., 2012. Efficient BackProp. Neural networks: Tricks of the trade. Springer Berlin Heidelberg. 9–48

Lewis-Peacock, J.A., Norman, K.A., 2013. Multi-voxel pattern analysis of fMRI data. Cogn. Neurosci. V.

Meier, T.B., Desphande, A.S., Vergun, S., Nair, V.A., Song, J., Biswal, B.B., Meyerand, M.E., Birn, R.M., Prabhakaran, V., 2012. Support vector machine classification and characterization of age-related reorganization of functional brain networks. Neuroimage 60, 601–613.

Misaki, M., Miyauchi, S., 2006. Application of artificial neural network to fMRI regression analysis. Neuroimage 29, 396–408.

Misaki, M., Kim, Y., Bandettini, P. A., Kriegeskorte, N., 2010. Comparison of multivariate classifiers and response normalizations for pattern-information fMRI. Neuroimage 53, 103–118. doi:10.1016/j.neuroimage.2010.05.051

Montavon, G., Lapuschkin, S., Binder, A., Samek, W., Müller, K., 2017. Explaining nonlinear classification decisions with deep Taylor decomposition. Pattern Recognit. 65, 211–222.

Mourão-Miranda, J., Bokde, A.L.W., Born, C., Hampel, H., Stetter, M., 2005. Classifying brain states and determining the discriminating activation patterns: Support Vector Machine on functional MRI data. Neuroimage 28, 980–995.

Mourão-Miranda, J., Friston, K.J., Brammer, M., 2007. Dynamic discrimination analysis: A spatial-temporal SVM. Neuroimage 36, 88–99.

Norman, K.A., Polyn, S.M., Detre, G.J., Haxby, J.V, 2006. Beyond mind-reading: multi-voxel pattern analysis of fMRI data. Trends Cogn. Sci. 10, 424–30.

Poldrack, R.A., Gorgolewski, K.J., 2017. OpenfMRI: Open sharing of task fMRI data. Neuroimage 144, 259–261.

Polyn, S.M., Natu, V.S., Cohen, J.D., Norman, K. A., 2005. Category-specific cortical activity precedes retrieval during memory search. Science 310, 1963–1966.

Power, J.D., Barnes, K.A., Snyder, A.Z., Schlaggar, B.L., Petersen, S.E., 2012. Spurious but systematic correlations in functional connectivity MRI networks arise from subject motion. Neuroimage 59, 2142–2154.

Rasmussen, P.M., Madsen, K.H., Lund, T.E., Hansen, L.K., 2011. Visualization of nonlinear kernel models in neuroimaging by sensitivity maps. Neuroimage 55, 1120–1131.

Ryali, S., Supekar, K., Abrams, D. A., Menon, V., 2010. Sparse logistic regression for whole-brain classification of fMRI data. Neuroimage 51, 752–764.

Saarimäki, H., Gotsopoulos, A., Jääskeläinen, I.P., Lampinen, J., Vuilleumier, P., Hari, R., Sams, M., Nummenmaa, L., 2016. Discrete Neural Signatures of Basic Emotions 2563–2573.

Schrouff, J., Phillips, C.L.M., 2012. Multivariate Pattern Recognition Analysis: Brain Decoding, in: Schnakers, C., Laureys, S. (Eds.), Coma and Disorders of Consciousness. Springer London, London, pp. 35–43.

Schrouff, J., Rosa, M.J., Rondina, J.M., Marquand, a. F., Chu, C., Ashburner, J., Phillips, C., Richiardi, J., Mourão-Miranda, J., 2013. PRoNTo: Pattern recognition for neuroimaging toolbox. Neuroinformatics 11, 319–337.

Smith, S. M., 2002. Fast robust automated brain extraction. Hum. Brain Mapp., 17, 143–55.

Smith, S.M., Jenkinson, M., Woolrich, M.W., Beckmann, C.F., Behrens, T.E.J., Johansen-Berg, H., Bannister, P.R., De Luca, M., Drobnjak, I., Flitney, D.E., Niazy, R.K., Saunders, J., Vickers, J., Zhang, Y., De Stefano, N., Brady, J.M., Matthews, P.M., 2004. Advances in functional and structural MR image analysis and implementation as FSL. Neuroimage 23, 208–219.

Stelzer, J., Chen, Y., Turner, R., 2013. Statistical inference and multiple testing correction in classification-based multi-voxel pattern analysis (MVPA): Random permutations and cluster size control. Neuroimage 65, 69–82.

Stelzer, J., Buschmann, T., Lohmann, G., Margulies, D.S., Trampel, R., Turner, R., 2014. Prioritizing spatial accuracy in high-resolution fMRI data using multivariate feature weight mapping. Front. Neurosci. 8, 1–8.

Sundermann, B., Herr, D., Schwindt, W., Pfleiderer, B., 2014. Multivariate classification of blood oxygen level-dependent fMRI data with diagnostic intention: A clinical perspective. Am. J. Neuroradiol. 39, 848–855.

Tettamanti, M., Rognoni, E., Cafiero, R., Costa, T., Galati, D., and Perani, D. (2012). Distinct pathways of neural coupling for different basic emotions. NeuroImage, 59, 1804–17.

Wang, Z., Peterson, B.S., 2008. Constrained least absolute deviation neural networks. IEEE Trans. Neural Networks 19, 273–283.

Woolrich, M. W., Jbabdi, S., Patenaude, B., Chappell, M., Makni, S., Behrens, T., Beckmann, C., Jenkinson, M., Smith, S.M., 2009. Bayesian analysis of neuroimaging data in FSL. NeuroImage, 45, S173–86.

Yamashita, O., Sato, M.A., Yoshioka, T., Tong, F., Kamitani, Y., 2008. Sparse estimation automatically selects voxels relevant for the decoding of fMRI activity patterns. Neuroimage 42, 1414–1429.

Zhao, H., Gallo, O., Frosio, I., Kautz, J., 2015. Loss Functions for Neural Networks for Image Processing 3, 1–11.

Åberg, M.B., Wessberg, J., 2008. An evolutionary approach to the identification of informative voxel clusters for brain state discrimination. IEEE J. Sel. Top. Signal Process. 2, 919–928.

